# Different aspects of social relationships contribute to subjective well-being via different functional connectomes

**DOI:** 10.1101/714618

**Authors:** Takashi Itahashi, Neda Kosibaty, Ryu-ichiro Hashimoto, Yuta Y. Aoki

## Abstract

The achievement of improved subjective well-being (SWB) is a worldwide issue. Prior studies using self-report questionnaires have demonstrated that better social relationships (SR) form the foundation for better SWB. To confirm the relationships with objective measure and investigate the biological underpinning, we conducted connectome-based prediction modeling with a 10-fold cross-validation, utilizing resting state functional magnetic resonance imaging data (*n* = 761). Two aspects of SWB (life satisfaction and positive affect) were successfully predicted using SR-related functional connections (FCs). The models also showed good prediction performance in a validation sample (*n* = 91), suggesting that our models are generalizable. All six aspects of SR considered were related to different sets of FCs with minimal overlap in edge level. The contributions of these FC sets to the two aspects of SWB were substantially different. In the future, effort should be made to improve all aspects of SR to achieve better SWB.

## Introduction

Since the 1980s, the topic of subjective well-being (SWB) has received a lot of attention from researchers, economists, and policy makers (Diener, 1984; Layard, 2010). Researchers established that good SWB are protective against the well-known impacts of major life events, and are related to a lower chance of physical and mental illness, as well as to increased longevity (Diener & Chan, 2011; Luhmann, Hofmann, Eid, & Lucas, 2012; Wood & Joseph, 2010). Economic research has shown that average SWB increase as the nation average income increases, contrary to the phenomenon known as “Easterlin Paradox” (Frank, 2012). However, a survey by the Organization for Economic Co-operation and Development revealed a lack of improvement in SWB in many of its member countries over the past decade, despite improvements in income and health (OECD, 2017).

SWB is a complex cognition which has global impacts. Researchers have examined how SWB consist and are associated with other social factors. One such factors is social relationships (SR). SR refer to a social structure made up of relationships with a wide range of people, including family members, neighbors, colleagues and peers, as well as emotional and instrumental supports (Goswami, 2011). SR have both positive aspects, such as friendship, emotional and instrumental supports, and negative aspects, such as perceptions of hostility, loneliness, and rejection by peers. Prior studies have reported that better SR are the foundation for better SWB (Siedlecki, Salthouse, Oishi, & Jeswani, 2014). However, such studies only utilized self-report questionnaires to examine the relationships, leaving potential objective and biological bases underlying the relationship between SR and SWB unexplored.

SWB have objective and biological correlates. Indeed, genetic research has shown that genetic factors account for about one third of SWB (Bartels, 2015). A recent genome-wide association study identified loci related to SWB (Okbay et al., 2016). Magnetic resonance imaging (MRI) studies revealed the brain regions associated with SWB (Davidson & McEwen, 2012; Takeuchi et al., 2014), and SR also involve specific brain regions. For example, both perceptions of hostility and rejection by peers are associated with functional connections (FCs) related to emotional processing (Masten et al., 2011; Moses-Kolko et al., 2010), while loneliness is associated with social brain regions (Kanai et al., 2012).

Given that relationships between SR and SWB were examined using only subjective measures, confirming these relationships using objective measures, and investigating the biological basis will deepen our understanding and provide a potential clue to improve SWB. Prior studies have shown that both SR and SWB are associated with FCs and that SR lead SWB (Davidson & McEwen, 2012; Masten et al., 2011; Siedlecki et al., 2014). Thus, we hypothesized that SR have their neural correlates which underlie SWB. To address this hypothesis, we analyzed resting-state functional MRI (R-fMRI) and phenotypic data collected as a part of the Human Connectome Project (Van Essen et al., 2013).

## Results

### Prediction of SWB by models using FCs associated with SR

After preprocessing the publicly available R-fMRI data, we obtained a 376 × 376 matrix based on Glasser’s atlas (Glasser et al., 2016), consisting of 360 cortical parcels and 16 subcortical parcels, as regions of interest (ROIs). To examine whether SR have neural correlates that underlie SWB, we used connectome-based prediction modeling (CPM) (Shen et al., 2017) with a 10-fold cross-validation (CV). In this framework, we extracted FCs associated with SR and then constructed models for predicting SWB using FCs associated SR. Fig. 1 shows an overview of our analytical procedures and detailed procedures are described in Materials and Methods. For clarity, we defined sets of FCs associated with SR as SR networks throughout this manuscript.

**Fig. 1.**
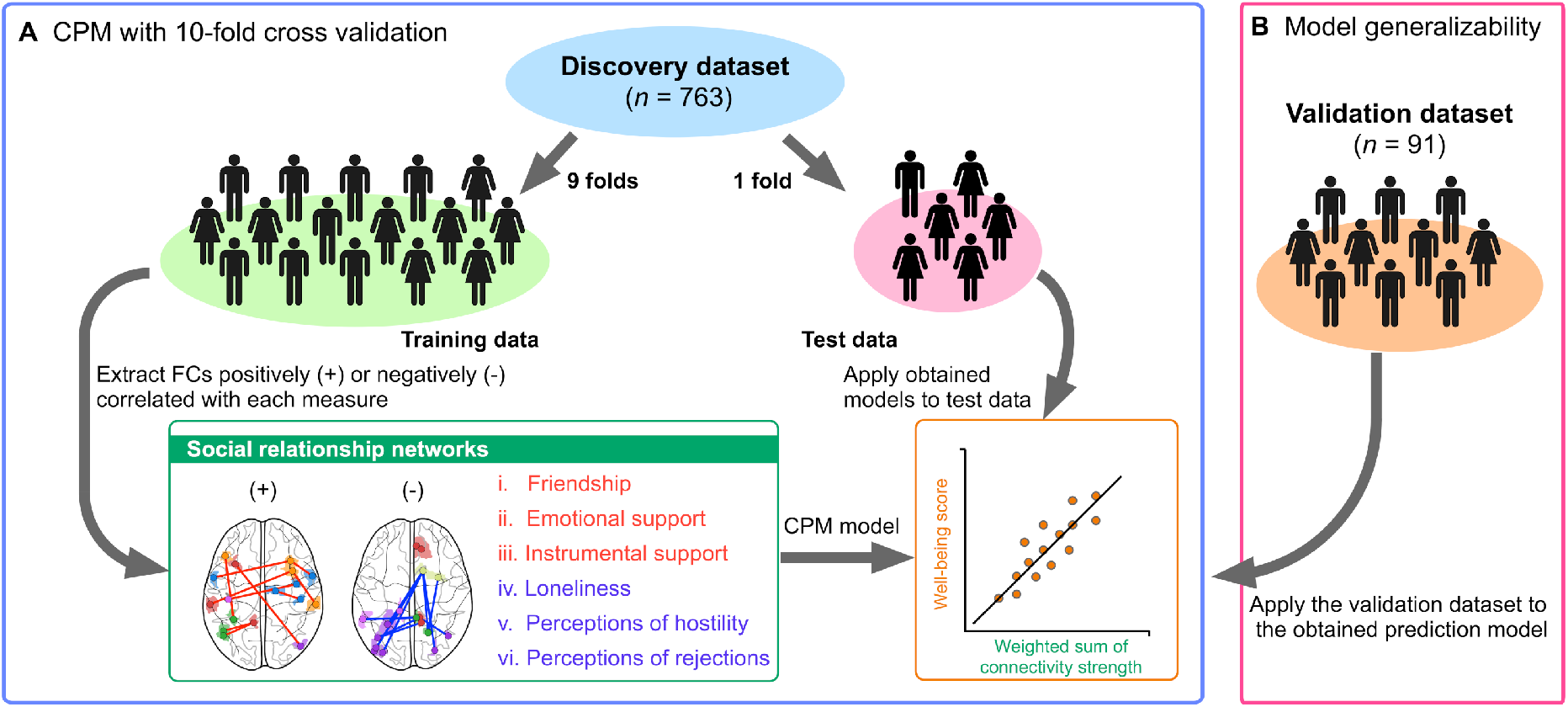
Schematic representation of the analytical procedures used in this study. Connectome-based predictive modeling (CPM) was performed using a 10-fold cross validation (CV) in the discovery dataset (*n* = 763). The number of subjects was divided into 10 folds. At each iteration, nine folds were used as a training data, to build models for predicting subjective well-being (SWB) via functional connections (FCs) associated with social relationships (SR). Pearson correlation coefficients were calculated between the strength of FCs and the degree of SR. The set of correlations was thresholded (*P* < 0.01) to create feature sets that were either positively (+) or negatively (-) correlated with SR. Then, the connectivity strength for each measure of SR was computed for each subject in the training data. The sets of FCs associated with SR were termed as “SR networks.” Multiple linear regression was used to build a model for predicting SWB from SR networks. The model was, then, applied to the left-out fold. The predictive power was assessed by the Pearson correlation coefficient between predicted and actual scores. Once statistically significant associations were observed, the model trained on the whole discovery dataset was applied to the validation dataset (*n* = 91), to evaluate the model generalizability.

Models with SR networks predicted two out of the three aspects of SWB: life satisfaction (*r* = 0.1202, 95%CI = [0.0438, 0.1927], *P* = 0.0009); and positive affect (*r* = 0.1086, 95%CI = [0.0409, 0.1808], *P* = 0.0027), as shown in Fig. 2A. These results remained when thresholds were set (*P* < 0.001, 0.005, and 0.05) for selecting relevant FCs (see Fig. S1).

**Fig. 2.**
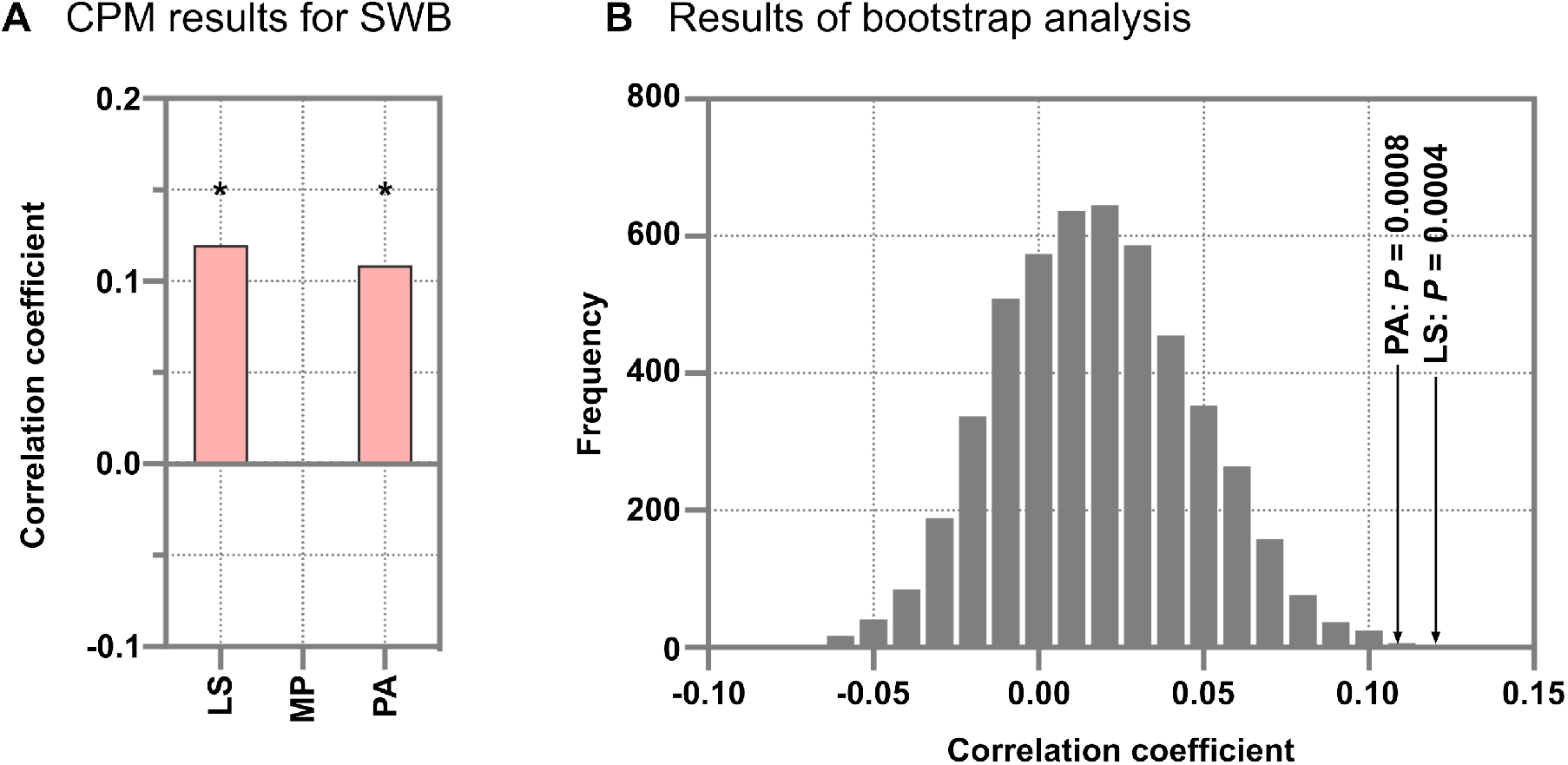
Performance of prediction models for subjective well-being using social relationship networks. (A) Bar graphs represent the prediction performances of the models for subjective well-being (SWB). The asterisk denotes *P* < 0.05 with Bonferroni correction. (B) The frequency of the different prediction performances is plotted in a bootstrap analysis with 5,000 iterations. At each iteration, the same number of functional connections (FCs) involved in social relationship (SR) networks was randomly selected from all 70,500 FCs except for the FCs involved in the SR networks in the 10-fold cross validation procedure. The highest correlation coefficients across all the null models were stored to construct the null distribution. This analysis indicated that the probabilities of prediction performances of life satisfaction *(r* = 0.1202) and positive affect (*r* = 0.1086) were small (life satisfaction: *P* = 0.0004 and positive affect: *P* = 0.0008), and demonstrated that SR networks were indeed associated with SWB. **Abbreviations**: CPM: connectome-based predictive modeling, LS: life satisfaction, MP: mean and purpose, and PA: positive affect.

### Bootstrap analysis to test the associations between SWB and SR networks

To examine whether SR networks were statistically significantly associated with SWB, bootstrap analysis was performed. Prediction performances were compared with null models that were built using randomly selected FCs not included in the pool of selected FCs in the actual models (Yahata et al., 2016).

As shown in Fig. 2B, models with SR networks remained statistically significant for life satisfaction (P = 0.0004) and positive affect (P = 0.0008), while mean and purpose did not reach statistical significance (P = 0.6948). These results demonstrated that the prediction performances of models with SR networks were indeed significant for the life satisfaction and positive affect of SWB.

### Generalizability of association between SWB and SR networks

Model generalizability was tested using the validation dataset from “100 Unrelated Subjects.” As shown in Fig. 3, the models demonstrated statistically significant prediction accuracies for both measures: life satisfaction: *r* = 0.3951, 95%CI = (0.2443, 0.5339), *P* = 0.0001; and positive affect: *r* = 0.4918, 95%CI = (0.3313, 0.6204), *P* < 0.0001. These results further support the association between the life satisfaction and positive affect of SWB, and SR networks.

**Fig. 3.**
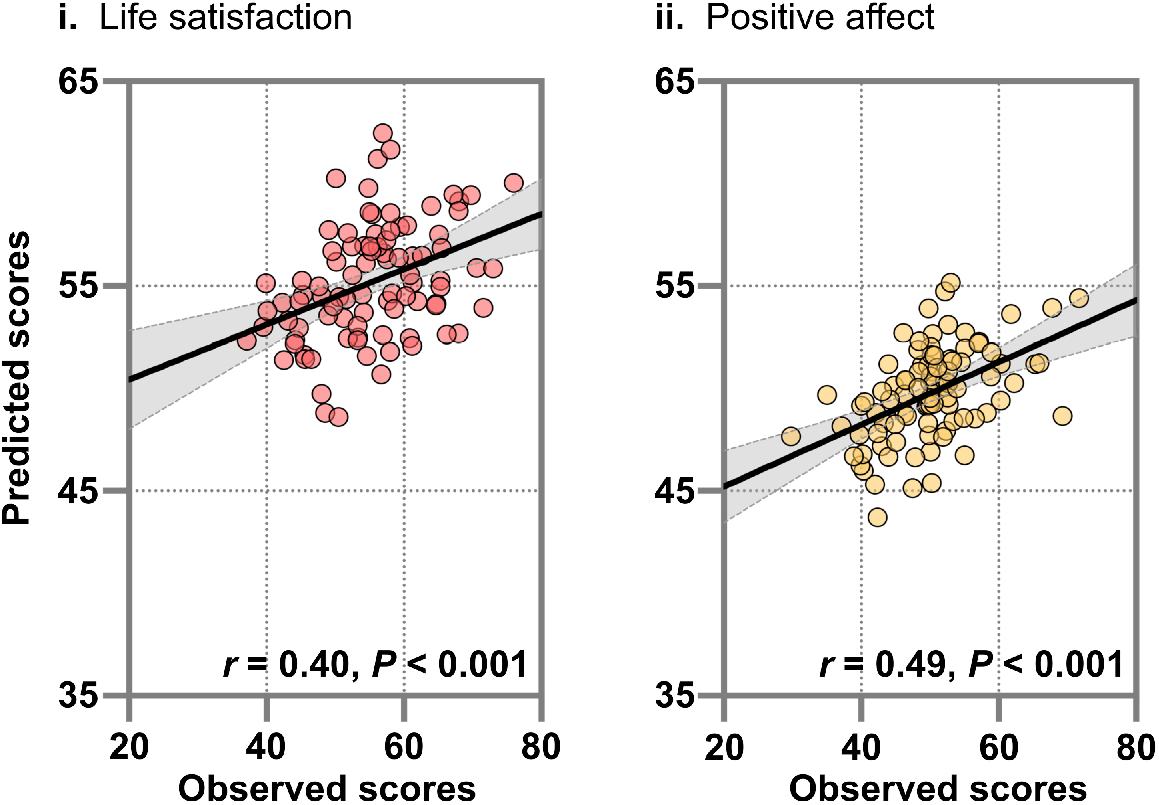
Generalizability of social relationship networks for predicting life satisfaction and positive affect. Scatter plots show the prediction accuracies of constructed models for *(i)* life satisfaction and *(ii)* positive affect in the independent validation dataset (*n* = 91). Models were constructed using all the discovery dataset (*n* = 763) with functional connections selected at least once during 10-fold cross validation. These models were then applied to the validation dataset. Prediction performance was estimated by computing the Pearson correlation coefficient between observed and predicted scores. The threshold for statistical significance was set to *P* <0.05.

### Reproducible associations between FCs and SWB and SR

To investigate whether the relationships of FCs with SWB and SR were reproducible across datasets, we performed mass univariate analyses. Pearson correlation coefficients were calculated between the strength of FCs and the degrees of SWB and SR for each dataset. As shown in Fig. S2, the effects of SWB and SR on FCs over the two datasets were positively correlated (all *r* > 0.32), suggesting that these effects are reproducible. Permutation tests with 5,000 iterations confirmed statistically significant positive correlations for SWB and SR (all *P* < 0.001).

We next investigated whether relationships of SWB-FC and SR-FC were correlated in both datasets. The Pearson correlation coefficients of SWB-FC and SR-FC were computed for each FC. As shown in Fig. S3 and Fig. S4, statistically significant associations between life satisfaction and positive affect of SWB and SR were observed in both datasets. These results reflect the similarities in the SWB-FC and SR-FC relationships.

### Associations between SWB and personality trait-related networks

In addition to SR, personality traits may have influences on the extent of SWB (Kong, Hu, Xue, Song, & Liu, 2015; Okbay et al., 2016; Strickhouser, Zell, & Krizan, 2017). To investigate whether personality traits were also associated with SWB, we built prediction models with 10-fold CV, replacing SR with personality traits. The models did not predict any of the measures of SWB (all *P* > 0.078, after Bonferroni correction; Fig. S5). These results suggest that SR, rather than personality traits, are associated with two aspects of SWB: life satisfaction and positive affect.

### Contributions of SR network to SWB

To examine which SR networks contributed significantly to the prediction of life satisfaction and positive affect, Wilcoxon signed rank tests were performed on the weight coefficients obtained during 10-fold CV. The threshold for statistical significance was set to *P* < 0.05, with Bonferroni correction for 12 comparisons.

Statistical analyses revealed that networks of positive SR, such as friendships and instrumental supports, mainly contributed to prediction of life satisfaction (Fig. 4A), while networks including aspects of loneliness, perceptions of hostility, and instrumental support dominantly contributed to prediction of positive affect (Fig. 4B). These results suggest that different SR networks are differently linked to life satisfaction and positive affect.

**Fig. 4.**
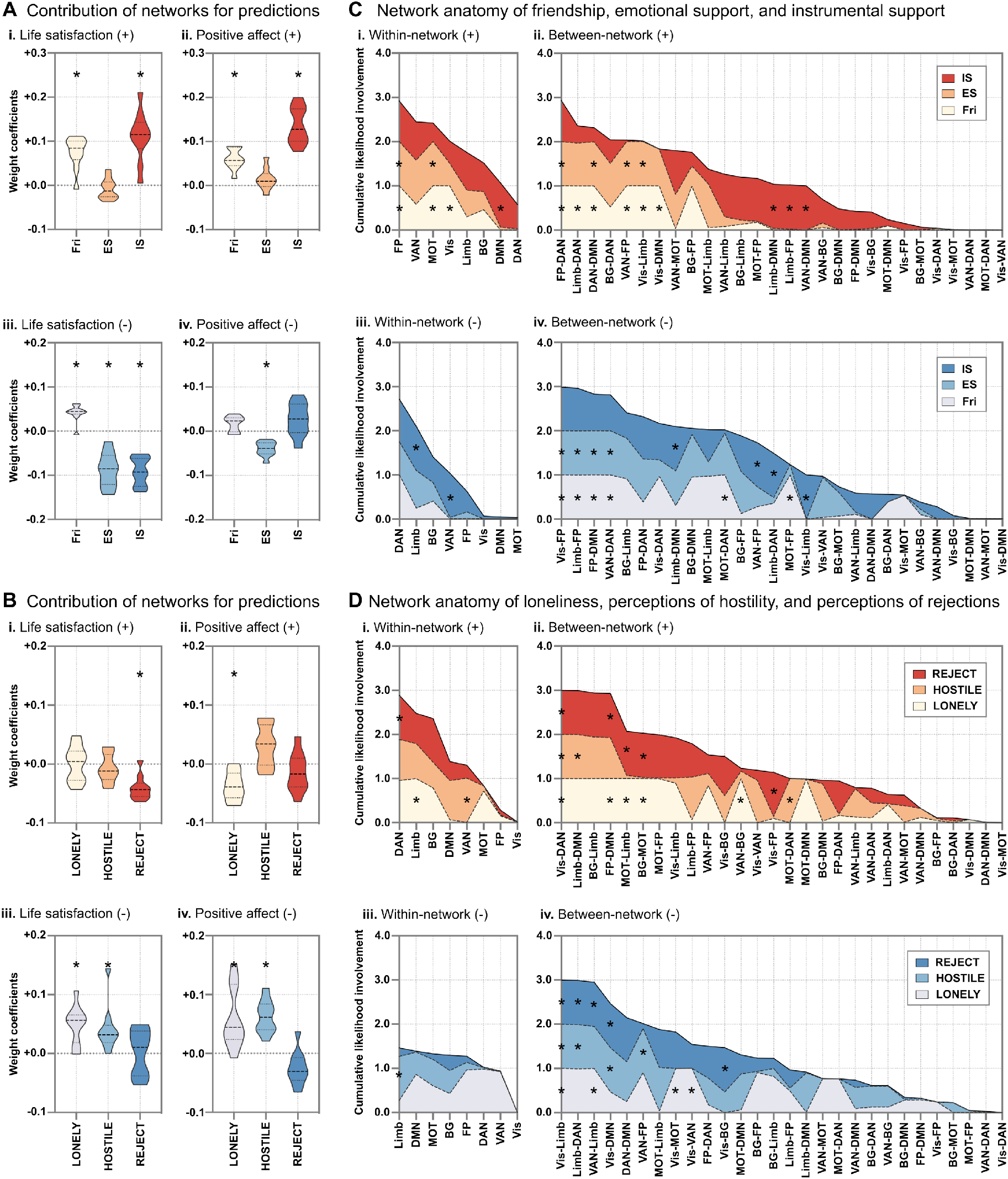
Anatomy of social relationships networks and contribution of these networks to the prediction of subjective well-being. Violin plots show the contribution of (A) positive and (B) negative networks to the prediction of life satisfaction (i and iii) and positive affect (ii and iv). The statistical significance of the contributions was assessed using the Wilcoxon signed rank tests. The asterisk represents the statistical significance after Bonferroni multiple comparison correction. For (C) positive and (D) negative social relationships, edge overlap (*i* and iii) within and (ii and iv) between eight a priori networks (Yeo et al., 2011) and our social relationship networks were plotted for sets of FCs (i and ii) positively and (iii and iv) negatively associated with each measure of social relationships. Each layer plot shows the sum of likelihood (1 – *P* value) estimated from the probability of edges shared between a priori networks and each social relationship networks. In all the plots, within-network and between-network pairs are sorted in the descending order. **Abbreviations**: BG: basal ganglia, DAN: dorsal attention network, DMN: default mode network, ES: emotional support, FP: frontoparietal, Fri: friendship, HOSTILE: perceptions of hostility, IS: instrumental support, Limb: limbic, LONELY: loneliness, MOT: somatomotor, REJECT: perceptions of rejections by peers, VAN: ventral attention network, and Vis: visual.

### Network anatomy of SR networks

Analyses were conducted to clarify the anatomy of SR networks. Binomial tests demonstrated that each aspect of SR was associated with around 0.7%–1.3% of FCs in the whole brain connections. In terms of positive aspects of SR (i.e., friendship, emotional support, and instrumental support), networks exhibited similar trends, except for the instrumental support network (Fig. 4C). For example, edges involved in the fronto-parietal (FP), somatomotor (MOT), and dorsal attention (DAN) networks were more likely to be involved in networks of friendship and emotional support. Edges stemming from limbic (Limb) and default mode (DMN) networks were likely to be associated with instrumental support. In contrast, edges from DMN and visual (Vis) networks were likely to be associated with negative aspects of SR (Fig. 4D). These results indicate that each of the SR networks was substantially different.

### Shared FCs within SR networks

Since we obtained 12 SR networks, our conjunction analyses were restricted. Conjunction analyses did not find any shared edges across the six SR networks for positive/positive, negative/negative, positive/negative, or negative/positive combinations. As shown in Fig. 5A and Table S3, networks of the positive aspects of SR (i.e., friendship, emotional support, and instrumental support) shared only 10 positive edges and 6 negatives edges. Positive edges consisted mainly of edges from the right insula and left precuneus to other regions, such as the bilateral precentral gyri and the left inferior and middle temporal gyri, while negative edges were from regions in DAN to other regions, such as the left inferior frontal gyrus and inferior parietal lobule. In contrast, networks of the negative aspects of SR (i.e., loneliness, perceptions of hostility, and perceptions of rejection by peers) contained eight positive edges mainly from the DMN and Limb, and six negative edges stemming from regions in the Vis to other regions involved in the DAN, DMN, and Limb (Fig. 5B and Table S3).

**Fig. 5.**
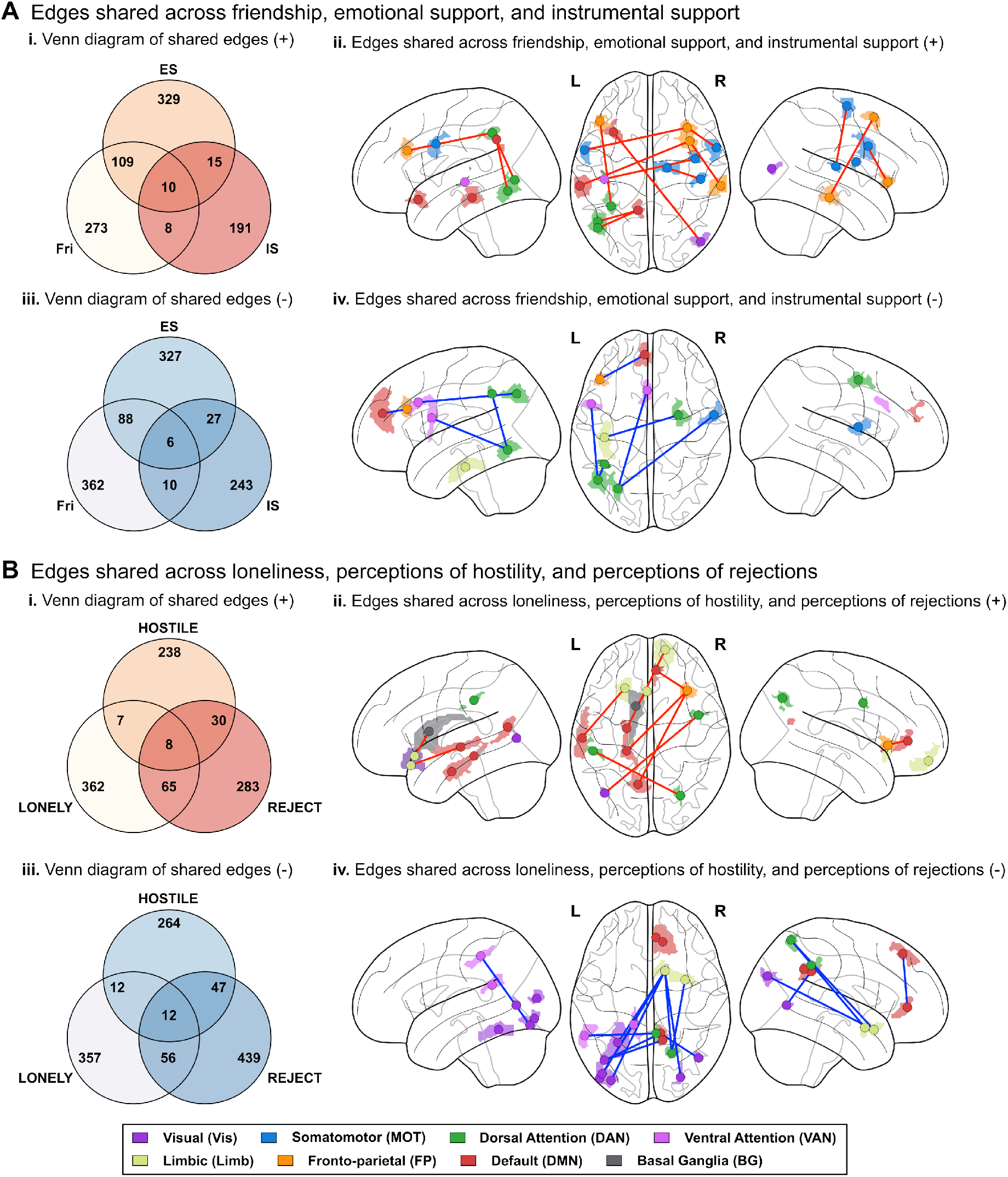
Shared functional connections (FCs) across (A) positive and (B) negative social relationships. Venn diagrams represent the number of shared FCs (i) positively or (iii) negatively associated with each of (A) positive or (B) negative social relationships. (ii and iv) Distributions of shared FCs among all the three measures are visualized. **Abbreviations**: ES: emotional support, Fri: friendship, HOSTILE: perceptions of hostility, IS: instrumental support, L: left, LONELY: loneliness, R: right, and REJECT: perception of rejection by peers.

## Discussion

The current study demonstrated that SR were associated with functional networks that were eventually related to two aspects of SWB: life satisfaction and positive affect. More specifically, life satisfaction was associated with all six aspects of SR, while positive affect was linked with all, except for perceptions of rejection by peers. Although all aspects of SR were connected to SWB, the degree of contribution of each differed. The neural bases were substantially different from each other in edge level. These FC-SWB relationships were not reflected in personality traits.

The R-fMRI data helped us disentangle the complexity of SWB. At the network level, friendship and emotional support showed similarity in FCs in both positive and negative directions. Given that the friendship and emotional support questionnaires ask about similar aspects of social relationships, the similarity in the network patterns is understandable. FCs negatively associated with friendship were positively associated with both life satisfaction and positive affect, while FCs negatively associated with emotional support were negatively related to life satisfaction and positive affect. These results may suggest that friendship may boost FCs among some regions, while lack of emotional support leads to anti-correlation of activity among some regions in the brain. Instrumental support had completely different FC networks from those of friendship and emotional support, while it had the biggest effect on both life satisfaction and positive affect. Although they were different, FCs negatively related to loneliness and perception of hostility showed positive influence on both life satisfaction and positive affect. FCs related to the perception of rejection by peers was negatively associated with life satisfaction. These SR networks hardly overlapped in edge level. Such differences in the way the SR-related FCs contributed to aspects of SWB and differences in edges that were related to each aspect of SR may contribute to the complexity of SR-SWB relationships. Given that all SR networks were shown to be related to at least one of the three aspects of SWB, the current findings suggest that improvement in any of the SR parameters may lead to improvements in SWB. These observations indicate that it is important to improve all the aspects of SR to achieve better SWB.

The results of this study were in line with the results of prior studies that examined the relationships between SR and SWB. However, meaning and purpose did not have any links with any of the SR networks. There are two potential explanations for this discrepancy within SWB. First, compared with life satisfaction and positive affect, meaning and purpose is a product of higher order cognition, and consists of more complex combination of cognitive components. Thus, SR networks were not related meaning and purpose. Another explanation is that meaning and purpose can be inward and own feeling, which is independent from SR. The lack of links between mean and purpose and SR networks suggests that this aspect of SWB should be addressed differently from the other two aspects.

The current findings have some limitations. First, we used a cross-sectional dataset to examine the triad associations between SWB, FC, and SR. Although we confirmed the main findings using a validation dataset, the causal relationship among them is still unclear. Future longitudinal investigations are necessary to elucidate the causal relationships. Second, although we demonstrated associations between the SR and SWB, the types of SR that influence on SWB may vary with developmental stages. For instance, relationships with parents might be a critical factor from infancy to early adolescence (Chen, Haines, Charlton, & VanderWeele, 2019; Itahashi et al., 2019), while the influence of relationships with peers or other factors may be pronounced in late adolescence to adulthood (Fortuin, van Geel, & Vedder, 2015). Future research with subjects from multiple life stages (Harms et al., 2018; Somerville et al., 2018) may help to further disentangle the complex associations between SWB and environmental factors.

The current study corroborated the relationships between SR and SWB. Each aspect of SR had a unique set of FCs, which was linked to at least one aspect of SWB. The current study suggests the necessity of improving all the aspects of SR to achieve better SWB.

## Materials and Methods (less than 3000 words)

### Human Connectome Project dataset

Data were collected as a part of the HCP (Van Essen et al., 2013). In this study, we used data from the “1200 Subjects” public data release (https://www.humanconnectome.org/study/hcp-young-adult) as a discovery dataset and those from the “100 Unrelated Subjects” set as a validation dataset. The S1200 release and 100 Unrelated Subjects set originally contained 1113 subjects and 100 subjects, respectively. Our analyses were restricted to 763 subjects (397 females and 366 males) and 91 subjects (51 females and 40 males) who completed the NIH toolbox (SR and psychological well-being) as well as personality traits, and who exhibited low head motion during R-fMRI scans (< 3 mm translation, < 3° rotations, and < 0.15 mm in mean FD).

The measures of psychological well-being obtained using the NIH toolbox, consisting of general life satisfaction, meaning and purpose, and positive affect (Salsman et al., 2014). We defined the psychological well-being as SWB because SWB usually include life satisfaction and positive affect although the definition varies across studies (Diener, Suh, Lucas, & Smith, 1999; Steptoe, Deaton, & Stone, 2015). SR evaluated in the project consisted of six measures: friendship; loneliness; perceptions of hostility; and perception of rejection by peers in daily social interactions; and emotional and instrumental support (Cyranowski et al., 2013). Personality traits were assessed using the NEO-Five Factor Inventory (NEO-FFI) (McCrae & Costa Jr, 2004). This contains 60 questions divided into five different personality domains with 12 questions for each, regarding: neuroticism; extraversion/introversion; openness to experience; agreeableness; and conscientiousness. The details of the phenotypes are shown in Table S1 and Table S2.

### R-fMRI data preprocessing and network construction

Minimally preprocessed R-fMRI data (Glasser et al., 2013) were obtained from the publicly available HCP database. Using an in-house MATLAB code, we applied additional processing procedures to the R-fMRI data, including the removal of the first 10 seconds of data for each run, and nuisance regression on the data. Nuisance regressors consisted of linear detrending, six head motion parameters, and averaged signals from the white matter, ventricles, and gray matter, as well as their derivatives. A band-pass filter (0.008 – 0.1 Hz) was then applied to the residuals. FD was calculated to identify motion-contaminated volumes (Power, Barnes, Snyder, Schlaggar, & Petersen, 2012). To reduce spurious changes in FCs due to subtle head motion during scans, a scrubbing method with 0.5 mm threshold of FD was applied, as described in a previous study (Power et al., 2012).

We used Glasser’s 376 surface-based brain regions (Glasser et al., 2016) as ROIs, to characterize an individual’s functional network. Pearson correlation coefficients were calculated among all possible pairs of ROIs, yielding a 376 × 376 FC matrix for each subject. Fisher’s r-to-z transform was further applied to each correlation coefficient. These procedures yielded 70,500 unique FCs, excluding the diagonal elements of the FC matrix. To facilitate the interpretation of our findings, we identified the anatomical names of ROIs and the name of resting-state networks (RSNs) using the automated anatomical labeling (AAL) (Tzourio-Mazoyer et al., 2002) and a previous study (Yeo et al., 2011). We also added basal ganglia (BG) network to the RSN labels.

### Connectome-based prediction modeling for SWB using FCs associated with SR

We assumed that, if SR have their neural correlates which in turn underlie SWB, models using FCs associated with FCs could predict the degree of the SWB. To test the association, we built CPMs (Shen et al., 2017) with a 10-fold CV, for predicting SWB using SR networks. Fig. 1 shows an overview of our analytical procedures.

In each fold, we regressed out the effects of nuisance covariates, including sex, head motion (mean FD), and the reconstruction version of fMRI (Dubois, Galdi, Paul, & Adolphs, 2018) from FCs, SWB, and SR. To identify FCs related to SR, Pearson correlation coefficients were calculated between FCs and each measure of SR. The correlation coefficients were thresholded at *P* < 0.01 (Rosenberg et al., 2016) and separated into a set of FCs positively associated with SR and a set of those negatively associated with SR.

As a summary measure, connectivity strength (Rubinov & Sporns, 2010) was used to characterize each participant’s degree of FCs positively or negatively associated with each measure of SR, resulting in 12 summary measures for SR; both positive and negative associations with each of the six sub-scales of SR. We called these measures as “SR networks” throughout this study. A multiple linear regression model with the SR networks was trained to predict each measure of SWB. The trained model was used to predict the degree of SWB in the left-out fold. Prediction performance was evaluated by calculating the Pearson correlation coefficient between predicted and actual scores. The threshold for statistical significance was set to *P* < 0.05 with Bonferroni correction (i.e., *P* < 0.05/3 = 0.0167).

### Bootstrap analysis to test associations SWB and SR networks

Although analyses with the 10-fold CV found statistically significant associations between SR, the brain, and parts of SWB (see **Results**), the possibility of “over-fitting” in the prediction models remained (Whelan & Garavan, 2014). To examine whether SR networks were indeed associated with SWB, bootstrap analysis was performed (Yahata et al., 2016), in which prediction performances were compared with null models that were built using randomly selected FCs not included in the pool of selected FCs in the actual models.

Using a bootstrap analysis with 5,000 iterations, the Pearson correlation coefficient of the actual prediction models was compared with the null distribution derived from the bootstrap procedure. Note that we selected a best prediction performance (i.e., highest correlation coefficient) across all the null models at each iteration. The threshold for statistical significance was set to *P* < 0.05.

### Generalizability of association SWB and SR networks

Although statistically significant associations were observed with 10-fold CV and bootstrap analysis (see **Results**), the generalizability of observed associations remained. We tested the generalizability of prediction models using the validation dataset (*n* = 91). We built a prediction model using all the discovery dataset, with FCs selected during the 10-fold CV. Summary measures were computed using FCs that were involved in SR networks at least one time during the 10-fold CV procedure. The threshold for statistical significance was set to *P* < 0.05.

### Characterization of the functional anatomy of edges

To characterize the extent to which FCs contribute to the prediction, we performed binomial tests (Yamagata et al., 2018). We selected a subset of FCs based on correlations with each measure of SR in each fold. The feature selection procedure selected around 0.7 to 1.3% of FCs in the whole brain connections across all the measures of SR in each of the 10 validation folds. We assumed a binomial distribution with the probability of being selected from the set of FCs. The threshold of statistical significance was set to *P* < 0.05 with Bonferroni correction for 70,500 FCs with six measures of SR.

To further improve the interpretability of our findings, network anatomy was determined with a hypergeometric cumulative density function (Lake et al., 2019). We computed the probability of *m* significant edges in *K* possible edges in *n* edges from a finite population of size *M.* The threshold of statistical significance was set to *P* < 0.05 with Bonferroni correction for 36 comparisons. We also investigated the presence of shared edges within SR networks by computing the element-wise product across networks.

## Acknowledgments

Data were provided by the Human Connectome Project, WU-Minn Consortium (Principal Investigators: David Van Essen and Kamil Ugurbil; 1U54MH091657) funded by the 16 NIH Institutes and Centers that support the NIH Blueprint for Neuroscience Research; and by the McDonnell Center for Systems Neuroscience at Washington University. This work was partially supported by the JSPS KAKENHI (16H06396 to RH, 19K03370 and 19H04883 to TI, and 18K15493 to YYA).

## Competing interests

All the authors declare that no competing interests exist.

---

**Table S1.**
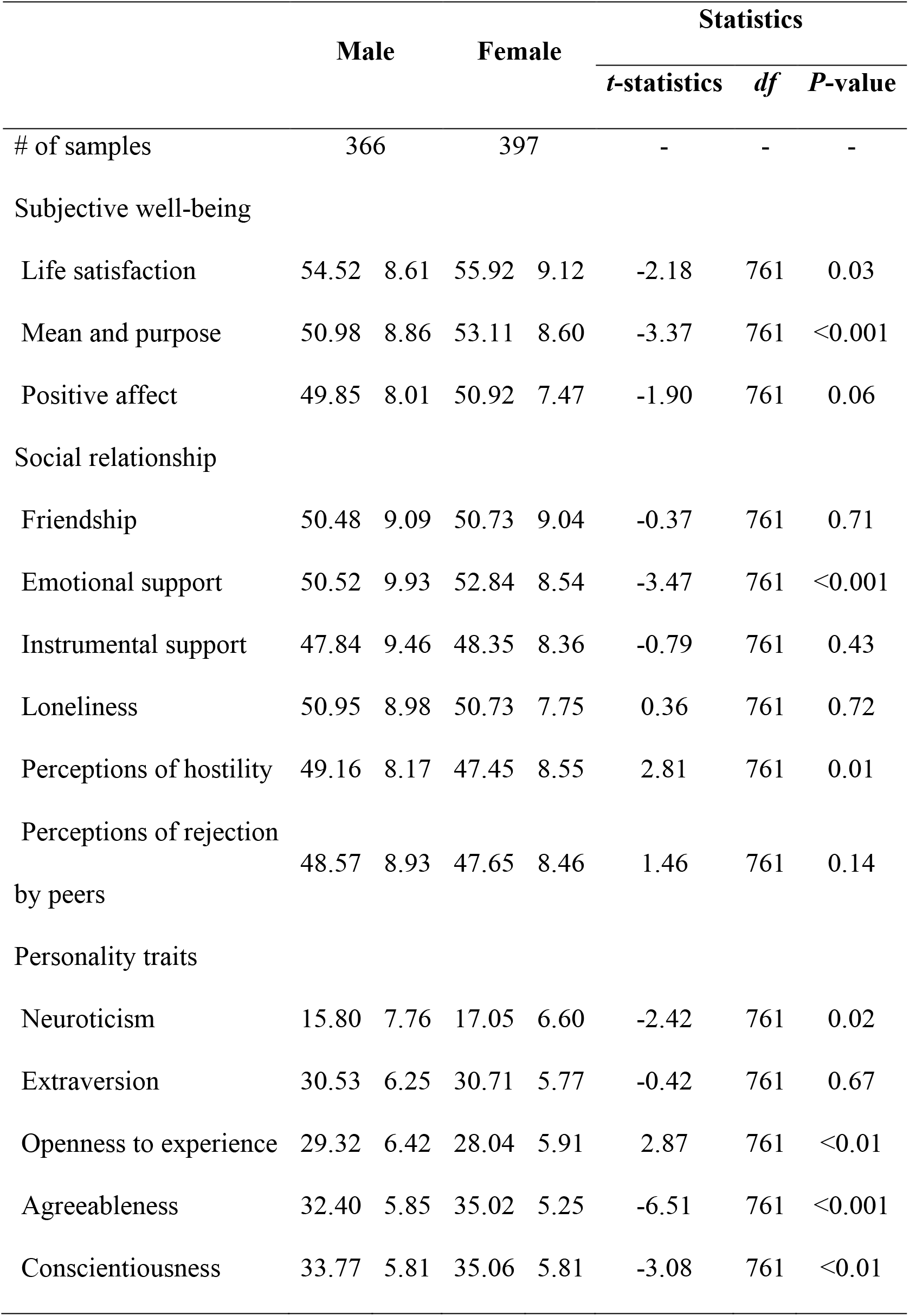
Phenotypic data of the discovery dataset

**Table S2.**
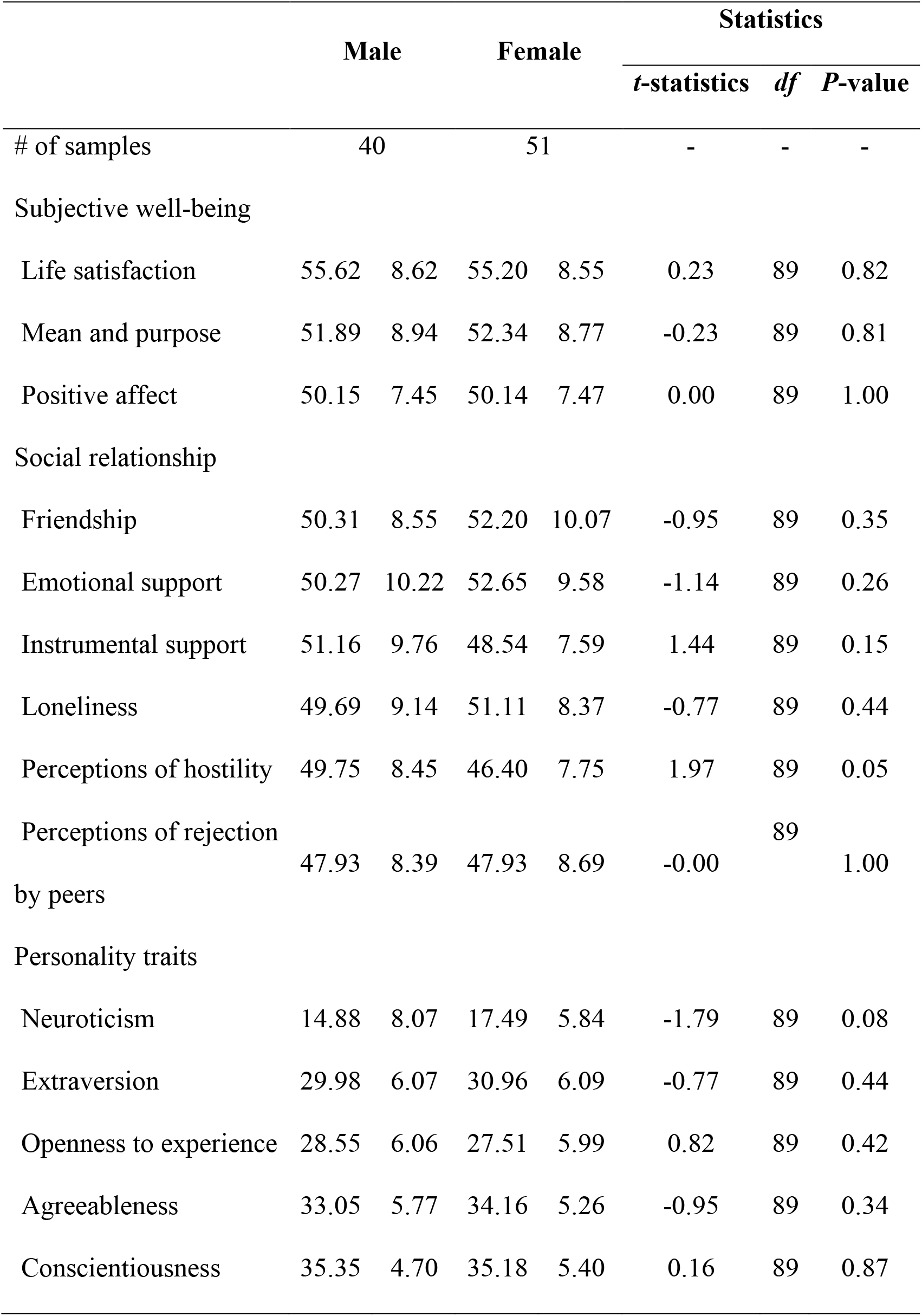
Phenotypic data of the validation dataset

**Table S3.** A list of functional connections (FCs) shared across social relationships.

| ROI1 |  |  | ROI2 |  |  | Type |
| --- | --- | --- | --- | --- | --- | --- |
| Glasser's area name | AAL Label | Network | Glasser's area name | AAL Label | Network |  |
| FCs shared across friendship, emotional support, and instrumental support |  |  |  |  |  |  |
| R.AVI | Insula_R | FP | L.6v | Precentral_L | MOT | + |
| L.AIP | Parietal_Inf_L | DAN | L.p9-46v | Frontal_Inf_Tri_L | FP | + |
| R.LO3 | Occipital_Mid_R | Vis | L.47s | Frontal_Inf_Orb_L | DMN | + |
| R.FOP2 | Rolandic_Oper_R | MOT | L.52 | Temporal_Sup_L | VAN | + |
| L.31pd | Precuneus_L | DMN | L.PH | Temporal_Inf_L | DAN | + |
| L.31pd | Precuneus_L | DMN | L.FST | Temporal_Mid_L | DAN | + |
| R.i6-8 | Frontal_Mid_R | FP | L.TE1m | Temporal_Mid_L | DMN | + |
| R.6mp | Supp_Motor_Area_R | MOT | R.A1 | Heschl_R | MOT | + |
| R.AVI | Insula_R | FP | R.6v | Precentral_R | MOT | + |
| R.TE1m | Temporal_Mid_R | FP | R.i6-8 | Frontal_Mid_R | FP | + |
| L.IP1 | Occipital_Mid_L | DAN | L.a24pr | Cingulum_Mid_L | VAN | - |
| L.p9-46v | Frontal_Inf_Tri_L | FP | L.9m | Frontal_Sup_Medial_L | DMN | - |
| L.PH | Temporal_Inf_L | DAN | L.6r | Frontal_Inf_Oper_L | VAN | - |
| R.6a | Frontal_Mid_R | DAN | L.TF | Temporal_Inf_L | Limb | - |
| L.IP2 | Parietal_Inf_L | FP | L.PH | Temporal_Inf_L | DAN | - |
| R.43 | Rolandic_Oper_R | MOT | L.IP1 | Occipital_Mid_L | DAN | - |
| FCs shared across loneliness, perception of hostility, and perception of rejection by peers |  |  |  |  |  |  |
| R.PEF | Precentral_R | DAN | L.MST | Occipital_Mid_L | Vis | + |
| R.AVI | Insula_R | FP | L.POS1 | Precuneus_L | DMN | + |
| L.A5 | Temporal_Mid_L | DMN | L.13l | Frontal_Inf_Orb_L | Limb | + |
| R.IPS1 | Occipital_Sup_R | DAN | L.PFt | Parietal_Inf_L | DAN | + |
| R.10pp | Frontal_Sup_Orb_R | Limb | L.EC | ParaHippocampal_L | DMN | + |
| R.AVI | Insula_R | FP | L.PreS | ParaHippocampal_L | DMN | + |
| Left caudate | Caudate_L | BG | L.25 | Olfactory_L | Limb | + |
| R.AVI | Insula_R | FP | R.a24 | Cingulum_Ant_R | DMN | + |
| R.31pv | Cingulum_Mid_R | DMN | L.MST | Occipital_Mid_L | Vis | - |
| R.31pd | Precuneus_R | DMN | L.MST | Occipital_Mid_L | Vis | - |
| R.pOFC | Olfactory_R | Limb | L.LO2 | Occipital_Inf_L | Vis | - |
| L.5mv | Cingulum_Mid_L | VAN | L.PIT | Fusiform_L | Vis | - |
| R.31a | Precuneus_R | FP | L.PSL | Temporal_Sup_L | VAN | - |
| R.pOFC | Olfactory_R | Limb | L.V3CD | Occipital_Mid_L | Vis | - |
| R.pOFC | Olfactory_R | Limb | L.VVC | - | Vis | - |
| R.31pv | Cingulum_Mid_R | DMN | R.MST | Temporal_Mid_R | Vis | - |
| R.pOFC | Olfactory_R | Limb | R.V7 | Occipital_Sup_R | Vis | - |
| R.Pir | Insula_R | Limb | R.VIP | Parietal_Sup_R | DAN | - |
| R.pOFC | Olfactory_R | Limb | R.VIP | Parietal_Sup_R | DAN | - |
| R.8BL | Frontal_Sup_R | DMN | R.a24 | Cingulum_Ant_R | DMN | - |

**Abbreviations:** L: left, R: right, ROI: region of interest
**Network:** BG: basal ganglia, DAN: dorsal attention network, DMN: default-mode network, FP: fronto-parietal, Limb: limbic, MOT: somatomotor, VAN: ventral attention network, and Vis: visual.

**Fig. S1.**
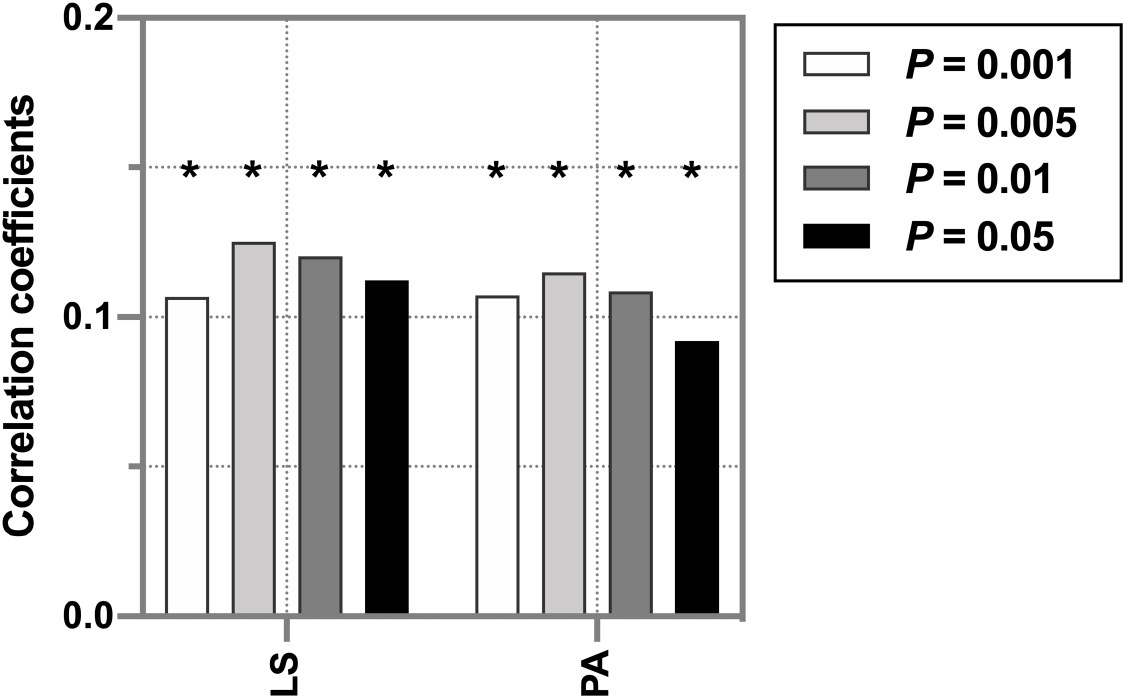
Performance of prediction models for subjective well-being using social relationships networks with different thresholds. Bar graphs represent prediction performances with different thresholds. Four different thresholds (P = 0.001, 0.005, 0.01, and 0.05) were used for selecting functional connections (FCs) associated with the social relationships. The asterisks denote *P* < 0.05. **Abbreviations**: LS: life satisfaction, and PA: positive affect.

**Fig. S2.**
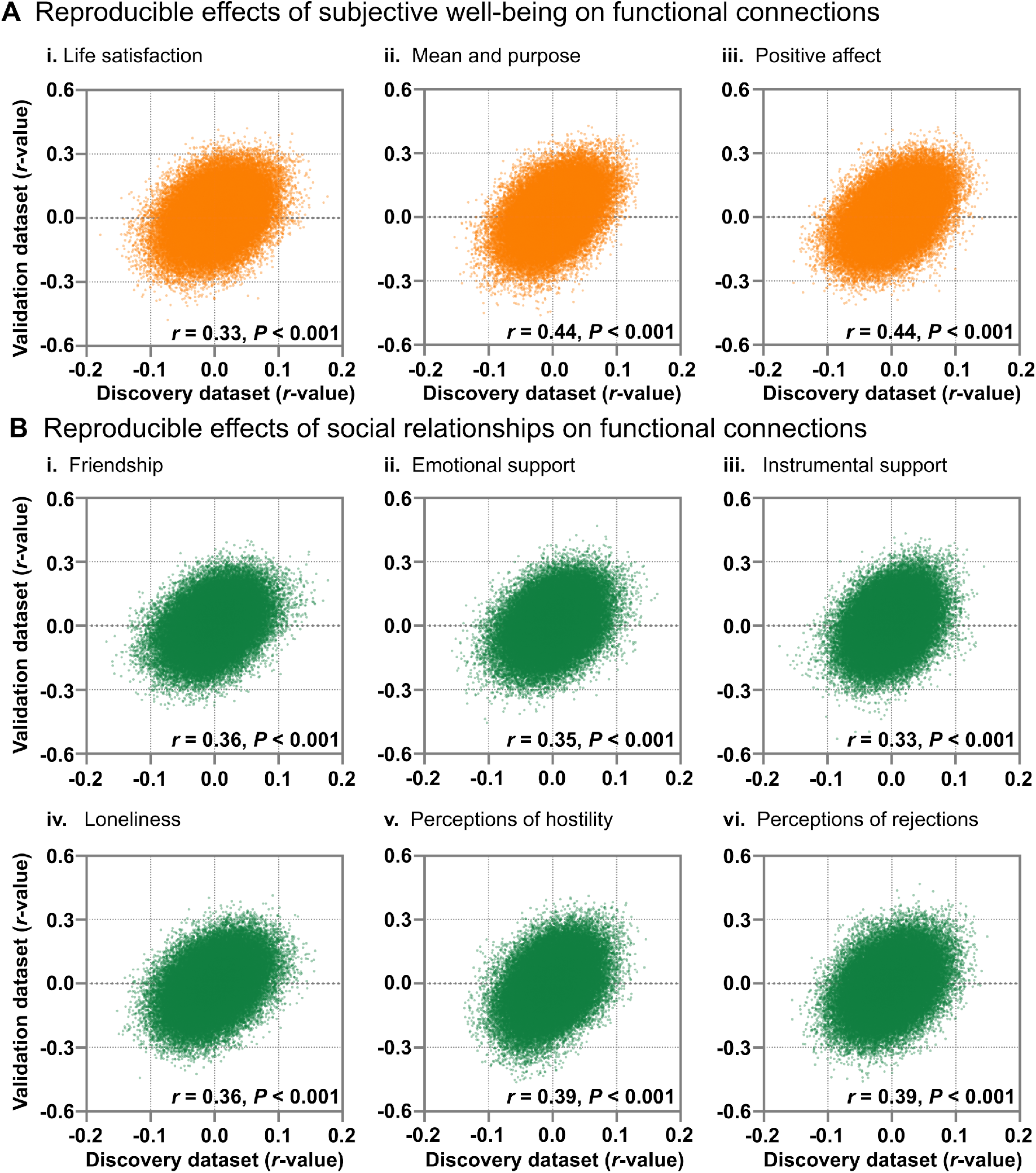
Reproducible effects of subjective well-being (A) and social relationships (B) across datasets. Scatter plots represent associations between the discovery dataset and the validation dataset. Permutation tests with 5,000 iterations confirmed that the effects of subjective well-being and social relationships were reproducible (all *P* < 0.001).

**Fig. S3.**
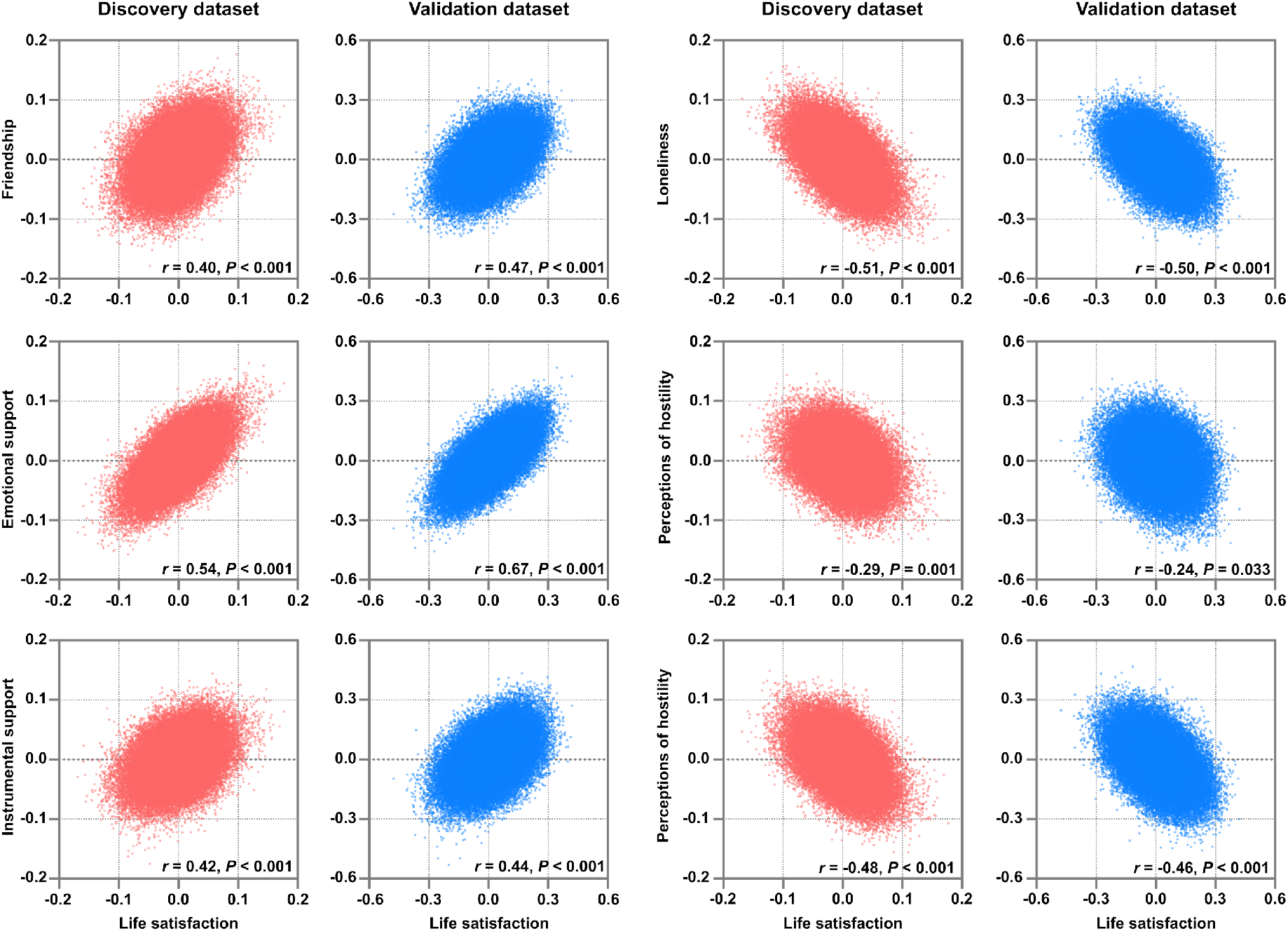
Similarities in effects of life satisfaction and social relationships on functional connections. Scatter plots represent associations between life satisfaction and social relationships on functional connections (FCs) separately for discovery (red) and validation (blue) datasets. P-values were estimated using permutation tests with 5,000 iterations. The threshold for statistical significance was set at *P* < 0.05.

**Fig. S4.**
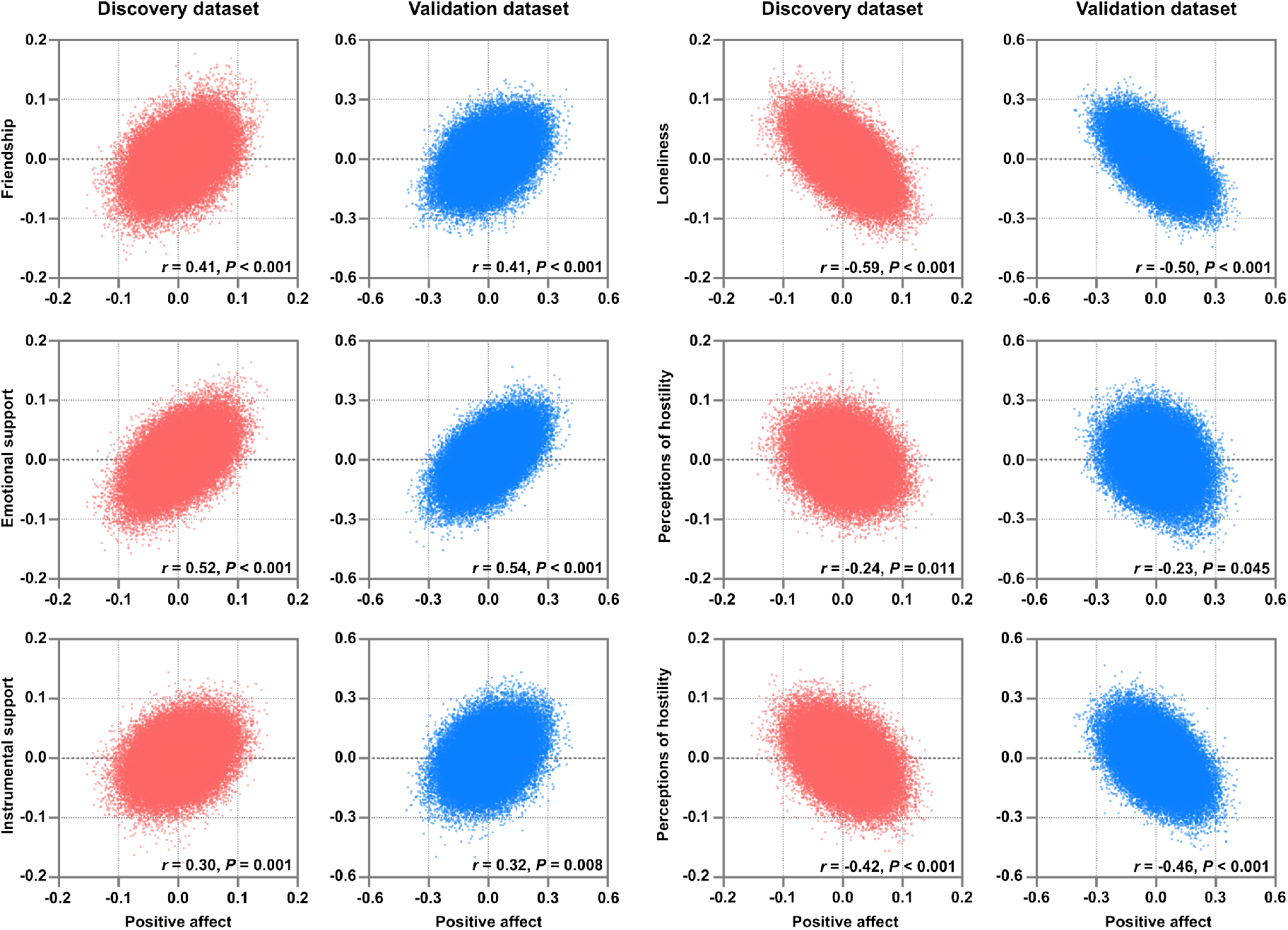
Similarity in effects of positive affect and social relationships on functional connections. Scatter plots represent associations between the effects of positive affect and those of social relationships on functional connections (FCs) separately for discovery (red) and validation (blue) datasets. *P*-values were estimated using permutation tests with 5,000 iterations. The threshold for statistical significance was set at *P* < 0.05.

**Fig. S5.**
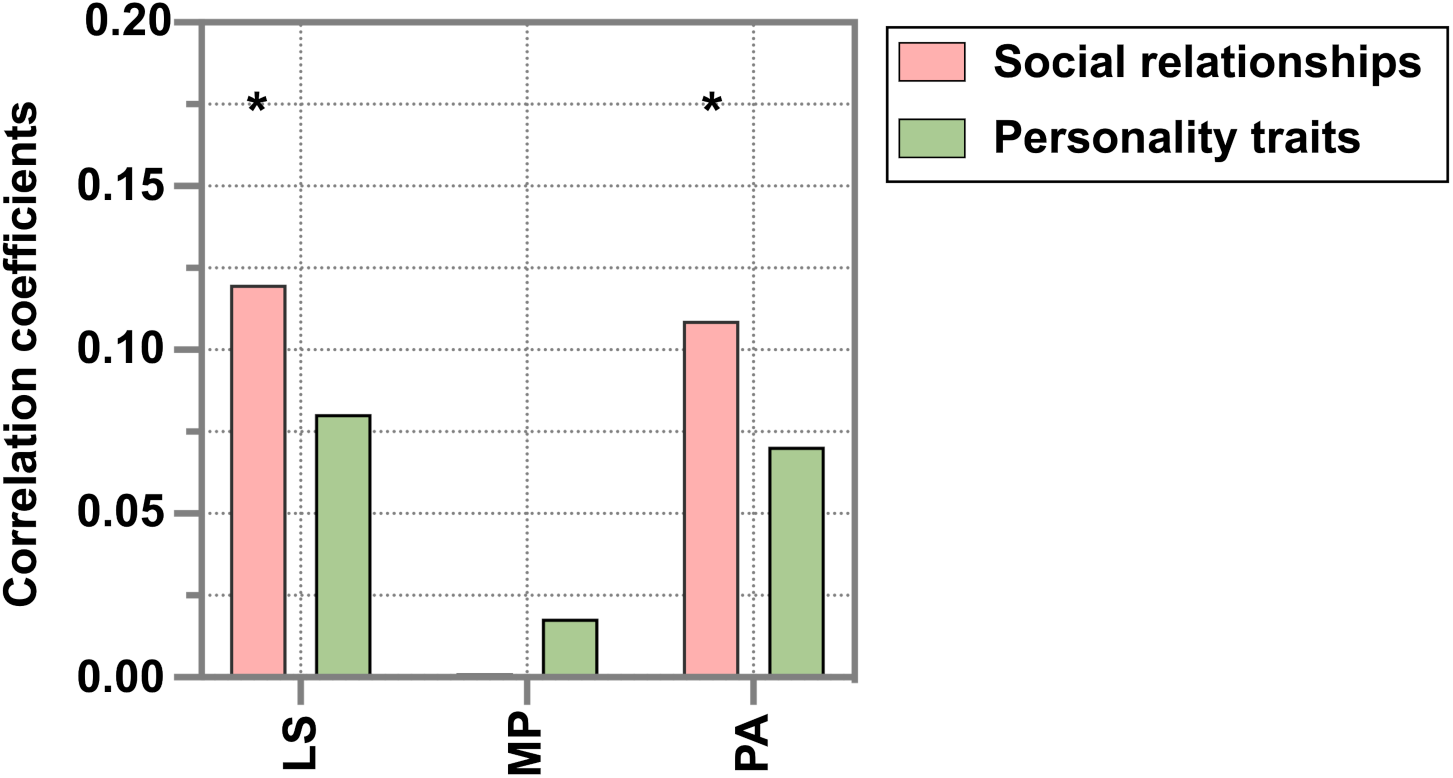
Performance of prediction models for subjective well-being using social relationships and personality traits. Bar graphs represent correlation coefficients between observed and predicted scores using social relationships (pink) and personality traits (green). The asterisks indicate statistical significance (i.e., *P* < 0.05/3)

